# Electrochemical Paper Fluidic Device for Extended Real-time Cocaine Monitoring in Simulated Sweat

**DOI:** 10.1101/2025.03.13.643163

**Authors:** Orlando S. Hoilett, Jenna F. Foust, Nikhil Davé, Bethany M. Balash, Dana L. Moryl, Anusha Kumar, Jacqueline C. Linnes

**Affiliations:** Weldon School of Biomedical Engineering, Purdue University, West Lafayette, IN 47907

## Abstract

Cocaine and other illicit substances are known to excrete from the body into sweat, making sweat an advantageous matrix for non-invasive detection and monitoring of substance use and abuse. However, existing detection of illicit substances in sweat remain single-use technologies that aren’t suitable for real-time, continuous monitoring over the course of an entire day. Long-term, continuous detection is hampered by large, bulky instrumentation, limited sensor stability for multi-hour measurements, and fluidic systems not optimized for minute and variable sweat flow rates. We demonstrate the continuous detection of 0.5 mM cocaine in simulated sweat via a paper fluidic device over the course of 17 hours. Our sensing system uses a biosensor functionalized with a cocaine-binding aptamer and a miniaturized electrochemical potentiostat. Our fluidics are driven by evaporation flow at physiological sweat rates (0.3-0.5 µL/min/cm^2^). Our development lays critical groundwork towards wearable, continuous detection of illicit substances in sweat.

## Introduction

Substance use disorder costs billions of dollars in increased healthcare, lost productivity, and crime [1]–[3]. Even with medical intervention, relapse rates among drug and alcohol dependence patients range from 40-60% within one year of starting treatment [4]–[7]. As a result, scientists, politicians, and public health officials alike are investigating using continuous on-body sensors for better treating substance use disorder, improving patient outcomes and quality of life [8]–[11].

Multiple methods, including immunoassays [12]–[14], mass spectrometry [15], fluorescence [16]–[18], and electrochemistry [19]–[21] have been used to detect cocaine and other illicit substances. Electrochemical techniques, in particular, provide simple, accurate, and sensitive detection [22]. As highly stable nucleic acid recognition elements, aptamers, and their use in electrochemical detection of cocaine have proven advantageous due to the method’s environmental stability and rapid response [21], [23], [24]. Electrochemical aptamer-based detection of cocaine has been conducted with several different biofluids such as blood serum [23], saliva [25], urine [19], [26] and even sewage [27]. While wearable, sweat-based sensors have not previously been used for real-time electrochemical cocaine detection, they have been used as sources for paper-based collection and off-body cocaine monitoring via mass spectrometry [15], [28]. Further, skin presents an attractive site for a continuous on-body monitoring, since sweat-based detection can be both non-invasive and discreet [14], [28], [29].

While paper has not been used for continuous cocaine detection in sweat, paper fluidic devices have been used for sweat collection and off-site analysis [14], [15], [28], [30]– [32]. Further, paper-fluidic devices have been designed for sweat collection on skin with real-time readouts of other analytes such as glucose [33], sodium [33], [34], alcohol [35], cortisol [36], and ammonium [37]. Paper provide ease-of-use, low cost, rapid analysis, easy disposal, portability, and low volume requirements, [38]–[43], making paper an appealing microfluidic platform for illicit substance detection at home and in the field [38], [40], [44]. While aptamers have been combined into paper microfluidic devices [45], continuous monitoring of aptamer responses under flow in paper has been elusive. This is due, in part, to the limited volume that a paper can hold, leading to saturation that stops flow past the sensor [46], [47], as well as the difficulty in ensuring the paper remains fully wetted at all times to prevent sensor degradation [48], [49]. The latter requirement is extremely critical to the success of paper-fluidic, aptamer-based sensors.

Here, we demonstrate the feasibility of a paper-based system for continuous, real-time electrochemical cocaine detection over the course of 17 hours. We modified a known cocaine-binding aptamer to incorporate methylene blue [21], a non-toxic electrochemically active redox tag, allowing for real-time analysis with accompanying instrumentation, our own custom-designed miniaturized potentiostat [50]. We then incorporated the sensor into a paper fluidic device that used evaporation to drive flow across the sensing electrodes [51] for the entire 17-hour monitoring period. We detail our results of detecting cocaine in a saline solution, in paper, and under flow at physiological sweat rates. The results demonstrate electrochemical analyte detection under flow over the course of an entire day in conditions that simulate continuous detection of cocaine in sweat.

## Materials and Methods

### Reagents, Materials, and Instruments

Ultrapure water was obtained from a Millipore Milli-Q system. Deionized water (DI) was obtained from the building’s in-house supply system. Phosphate-buffered saline (PBS) was purchased from MilliporeSigma (Part Number: P-5368, St. Louis, MO, USA) and dissolved in ultrapure water. PBS was used as the working buffer throughout the experiments and was kept at pH 7.4 and room temperature during use. Sulfuric acid (H_2_SO_4_) was obtained from MilliporeSigma (Part Number: 258105-100ML, St. Louis, MO, USA) and dissolved in appropriate volume of ultrapure water to obtain a 0.5 M and 0.05 M solution.

The cocaine-binding aptamer (5’ HSC11-AGACAAGGAAAATCCTTCAATGAAGTGGG TCG-MB 3’) was synthesized by LGC Biosearch Technologies (Petaluma, CA, USA). The aptamer includes a methylene blue (MB) redox tag on the 5’ end, making the aptamer electroactive, and a thiol group (HSC11) on the 3’ end, allowing the aptamer to self-assemble onto a gold electrode. The aptamer was provided by LGC as a lyophilized pellet and was dissolved in appropriate volume of ultrapure water upon receipt to create a 200 *μ*M aptamer solution. The aptamer solution was stored in the dark at 4^°^C until use.

Cocaine hydrochloride (cocaine) was purchased from MilliporeSigma (Part Number: C5776-1G, St. Louis, MO, USA) and stored securely in accordance with state and federal guidelines for controlled substances. 1-3 mg of cocaine was dissolved in PBS at the necessary concentrations. Cocaine solutions were made fresh for each experiment and disposed of in accordance with state and federal regulations.

Voltammetry was performed using a VSP-300 (Bio-Logic Science Instruments, Seyssinet-Pariset, France) as well as our own custom-designed miniaturized potentiostat, KickStat [50], as indicated. We used a gold (Au) working electrode (Part Number: MF-2014, Bioanalytical Systems, West Lafayette, IN, USA), silver silver-chloride (Ag/AgCl) reference electrode (Part Number: CHI111, CH Instruments, Austin, TX, USA), and a platinum (Pt) counter electrode (Part Number: CHI129, CH Instruments, Austin, TX, USA). Gold electrode polishing materials were purchase from CH Instruments, Inc. (Part Number: CHI120, Austin, TX, USA). The Ag/AgCl electrode was used as a “pseudo-reference” by removing the silver wire from its 1 M potassium chloride (KCl, Part Number: 3040-01, J.T. Baker) storage solution during experiments. Using the electrode as a “pseudo-reference” helped avoid cross-contamination between different solutions. We found that solutions diffused into and out of the storage solution and confounded earlier preliminary experiments.

A syringe pump was used (Model Number: Legato 210, KD Scientific Inc., Holliston, MA, USA) to model sweat rates in the multi-hour cocaine measurements. The paper fluidic device was made with Chromatography paper (Chr1, CAT No. 3001-861, GE Healthcare Life Sciences, Buckinghamshire, UK) and cut using a Silhouette Cameo cutter (Silhouette America, Inc., Lindon, Utah, USA). SelfSeal (Manufacturer: GBC, Germany), a pressure sensitive adhesive, was used to limit the evaporation of water from the paper fluidic device. A digital indoor temperature and humidity monitor (AcuRite 00613 Digital Hygrometer & Indoor Thermometer Pre-Calibrated Humidity Gauge) was used to record the temperature and humidity of the lab environment.

### Preparing Cocaine-Binding Aptamer-Modified Electrode

We followed the protocol from Hoilett *et al*. 2020 [50] to functionalize the working electrodes with cocaine aptamer. Briefly, the electrodes were chemically and physically cleaned with alumina polish, rinsed with DI water, then electrochemically cleaned with 0.5 M and 0.05 M H_2_SO_4_. The electrodes were incubated with a 400 nM aptamer solution in the dark and at room temperature for 2 hours, then in a 2 mM MCH solution in the dark and at room temperature for another 2 hours. Afterwards, the electrodes were gently rinsed with DI water for 10-15 seconds, then placed in PBS (pH 7.4 and room temperature) for 30 minutes before beginning electrochemical measurements. Incubating the electrode in PBS allowed molecules that were poorly immobilized onto the electrode surface to slowly unbind, stabilizing the functionalization and passivation layers [48], [52].

The prepared aptamer-modified electrode was analyzed using CV (7 cycles, -40 mV start and stop potential, from -0.45 V to -20 mV, 1 mV steps, at 50 mV/s) in cocaine-free PBS (pH 7.4 and at room temperature) to determine if the functionalization was successful.

### Characterizing Electrochemical Response in Chromatography Paper

To assess the effect of the chromatography paper substrate on the diffusion-controlled electrochemical response, we ran cyclic voltammetry (CV) (4 cycles, -0.04 V to -0.45 V, 1 mV step potential, at 25 mV/s, 50 mV/s, 100 mV/s, 200 mV/s) in a 5 mM potassium ferricyanide solution in a traditional electrochemical cell and in a saturated 0.5 cm x 2 cm strip of chromatography paper. We then plotted the anodic current vs. the square root of the scan rate and assessed the linearity of the responses using a simple linear regression in Prism 8. Solutions were kept at room temperature, a pH of 7.4, and referenced to an Ag/AgCl electrode.

#### Cocaine Measurements

Square wave voltammetry (SWV) was used to quantify the sensor response to 0 µM, 1 µM, 5 µM, 10 µM, 40 µM, 80 µM, 100 µM, and 180 µM of cocaine averaged across 4 identically prepared aptamer-modified electrodes. We used cocaine-free PBS buffer as 0 µM at the start of the experiment as well as before each subsequent cocaine concentration. Percent increase in signal was relative to the first measurement in PBS at the start of the experiment. Measurements of PBS before each subsequent measurement of cocaine were also measured relative to the initial measurement in PBS to quantify sensor drift. SWV was performed in a potential range of -0.05 V to -0.5 V (reduction scan), with a step potential of 1 mV, amplitude of 50 mV, and at a frequency of 62.5 Hz consistent with our previous work [50]. Solutions were kept at room temperature, a pH of 7.4, and referenced to an Ag/AgCl electrode.

Faradaic currents are usually determined by extrapolating a tangent to the baseline of the region preceding the reduction potential, then computing the peak currents relative to the tangent line. However, the preceding baseline of the SWV scans were often poorly defined (not flat) after repeated measurements. As a result, the peak currents were determined by subtracting the current at the reduction potential, usually near -0.33 V, from the current at the voltammogram’s next local minima, generally near -0.375 V.

### Determining Evaporation Rate of Water Through Chromatography Paper in Normal Lab Conditions

Experiments were conducted to determine the effect of humidity of the lab environment on the evaporation rate of water from the chromatography paper. Five (5), 2 cm x 2 cm pieces of chromatography paper were cut using a Silhouette Cameo cutter. Each piece of chromatography paper was weighed and then submerged in a 50 mL conical tube filled with 30 mL of ultrapure water for at least 30 minutes. Afterwards, each square was removed from the water. Excess water pooled on top of the paper was gently tapped off and the paper was weighed. The humidity of the room was also recorded using a commercial indoor temperature and humidity monitor. Each piece of paper was then placed on individual pieces of non-absorbent weighing paper in an open, undisturbed area of the lab and then weighed again after 30 minutes. The humidity of the room was recorded again. The evaporation rate in µL/min/cm^2^ as determined by dividing the difference in weights by 30 minutes and by 4 cm^2^ (the area of the paper).

### Determining Absorption Capacity of Chromatography Paper

Five 2 cm x 2 cm pieces of chromatography paper were cut using a Silhouette Cameo cutter. Each piece of chromatography paper was weighed while dry then submerged in 30 mL of ultrapure water for at least 30 minutes. The absorbance capacity of the paper was calculated by finding the difference between the paper’s dry weight and weight after being fully submerged in water. The difference in weights were then divided by the top surface area of the paper (2 cm x 2 cm).

### Multi-Hour Cocaine Measurements in Paper Fluidic Device

The paper fluidic device depicted in Figures 1A and 1B was cut using a Silhouette Cameo cutter. The device was first saturated with 0.5 mM cocaine by pipetting 162 *μ*L of cocaine evenly onto the paper fluidic device. The electrodes were placed onto the device as indicated in Figures 1B, 1C, and 1D. To simulate sweating, PBS and a 0.5 mM cocaine solution were sequentially delivered to the device using a syringe pump at flow rates ranging from 0.3-0.5 *μ*L/min/cm^2^ based on the humidity of the laboratory environment. Flow through the paper device was driven by evaporation by leaving the large 2 cm x 2 cm area (Figure 1A) uncovered. Care was taken to limit evaporation to only the large 2 cm x 2 cm area by covering the remaining portion of the device with SelfSeal. SWV (reduction scan, in a potential range of -0.05 V to -0.5 V, with a step potential of 1 mV, amplitude of 50 mV, and at a frequency of 62.5 Hz) was run every 10 minutes to monitor changes in current with the changing solution composition in the paper fluidic device. Solutions were kept at room temperature, a pH of 7.4, and referenced to an Ag/AgCl electrode.

**Figure 1.**
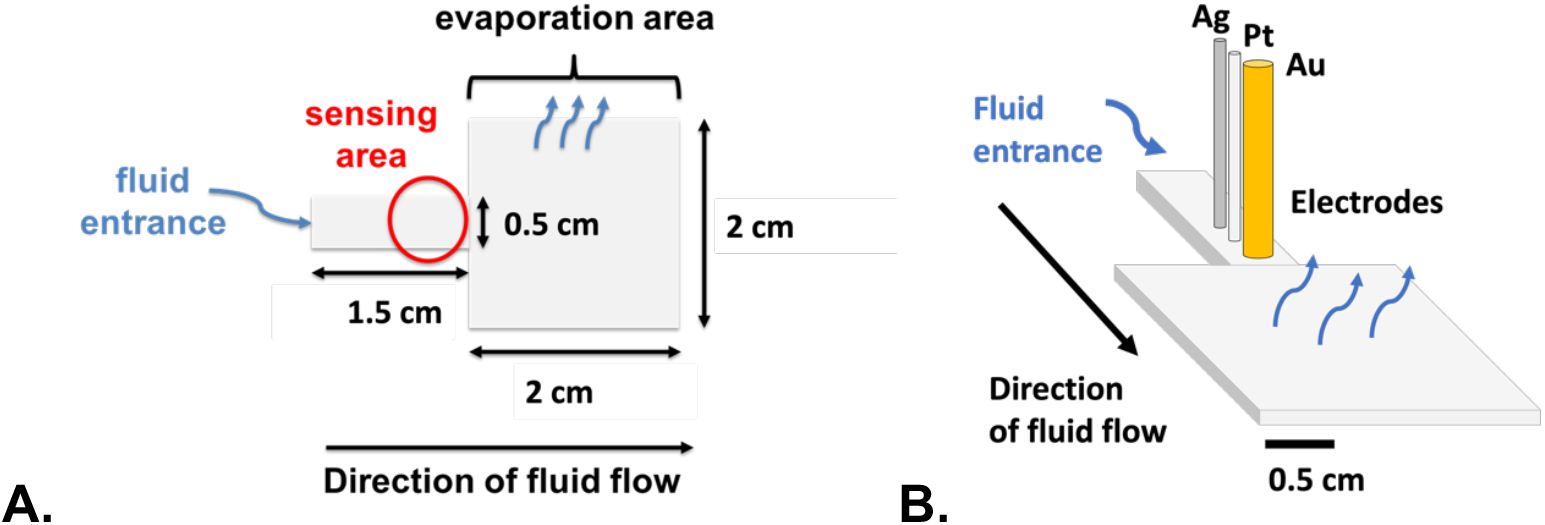
Paper fluidic device. **(A)** and **(B)** schematic of the paper fluidic device highlighting relevant dimensions and functional locations.

### Multi-Hour Detection of Cocaine in Paper Fluidic Device Using KickStat

We have previously developed a coin-sized potentiostat, called KickStat [50], for high-resolution electrochemical measurements. KickStat was programmed to run 20 consecutive SWV (reduction scan, in a potential range of -0.05 V to -0.5 V, with a step potential of 1 mV, amplitude of 50 mV, and at a frequency of 62.5 Hz) scans every 10 minutes throughout the duration of the experiment. The 20 scans were averaged to reduce variance. KickStat then determined the peak current from the averaged scan and compared the peak current to the next set of measurements performed 10 minutes later.

## Results and Discussion

### Cocaine Limit of Detection in Traditional Electrochemical Cell

Initial cocaine measurements in the traditional electrochemical cell were performed to ensure that the functionalization was successful. Figure 2 shows an example cyclic voltammogram of a fully prepared aptasensor in cocaine-free PBS in the traditional electrochemical cell. The redox peaks (oxidation on top and reduction on the bottom) have a near zero separation (∼20 mV), as expected for an adsorbed species with a reversible electrochemical reaction [53], indicating the functionalization was successful.

**Figure 2.**
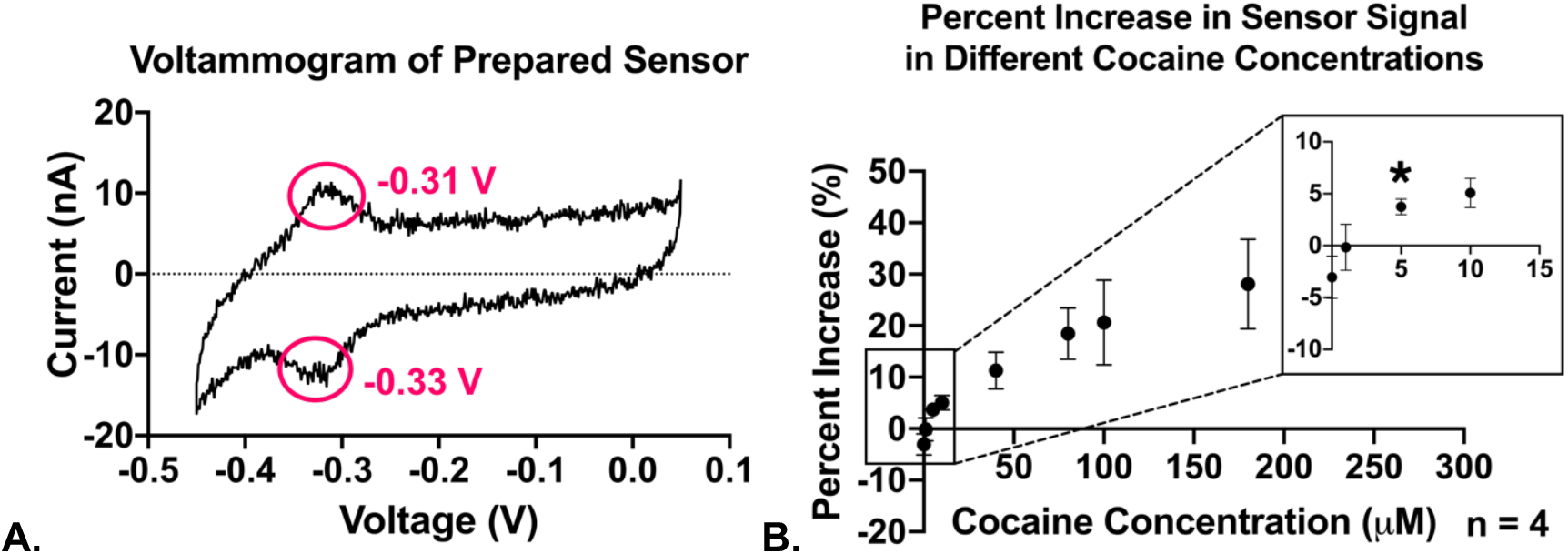
Cocaine sensing in a traditional electrochemical cell. (**A)** shows a cyclic voltammogram of a fully prepared aptasensor in PBS. **(B)** is the concentration-dependent signal gain of the sensor and inset detailing concentrations between 0 to 10 μM. The limit of detection is indicated with a * at 5 µM cocaine concentration (n = 4, error bars represent standard deviation).

To evaluate the limit of detection of our sensor, we measured the response of our cocaine aptamer to a range of concentrations of cocaine from 0 mM to 180 mM (Figure 2B). The percent increase in current measured for 5 µM cocaine (3.7% +/-0.76%) is greater than 1.645 standard deviations above the percent increase in current measured in the blank solution (−3.0% +/-2.0%). Therefore, following the calculations of Armbruster and Pry [54], we concluded that our sensor has a limit of detection of 5 µM, agreeing with others who reported detection limits less than 10 µM cocaine [21]. The negative control measurements with 0 µM cocaine (i.e., cocaine-free PBS) showed a negative percent increase relative to the baseline, also cocaine-free PBS, indicating a baseline drift of around 4-8% over the course 1 hour. Such baseline drift is expected with electrochemical measurement techniques [55]–[57]. Interestingly though, we observed a lower amount of drift (≤8%), after 1 hour of measurements, than others who saw drifts up to 35% using a similar aptamer-based biosensor [56]. This reduced drift may indicate a more well-developed passivation layer in our system. The passivation layer decreases the capacitive current of the electrode surface and has been shown to improve the long-term viability of aptamer-based biosensors [48], [49], [52]. Additionally, our aptamer is anchored to the electrode using an 11-carbon thiol which has shown greater stability, and may have contributed to our more stable baseline, than aptasensors using a 6-carbon thiol chain used by other groups [49], [56].

### Characterizing Electrochemistry in Chromatography Paper

Potassium ferricyanide is a well-defined Nernstian chemical [58]–[60], making it an optimal choice for characterizing chromatography paper as a platform for electrochemical analysis. The electrochemical response of potassium ferricyanide in unstirred solutions is expected to follow a diffusion-limited reaction, resulting in a linear dependence of the current to the square root of the scan rate, *ν*^1/2^ [53], [59]. Figure 3A depicts this linear dependence of current on the square root of scan rate in chromatography paper (R^2^ = 0.9987) and in the traditional electrochemical cell (R^2^ = 0.9995), indicating that paper does not adversely affect the diffusion of the electroactive species to the electrode surface. There is a smaller slope in the paper, 1.45 *μ*A/*ν*^1/2^, compared to in the traditional electrochemical cell, 2.10 *μ*A/*ν*^1/2^, indicating a small decrease in charge transfer efficiency. However, the decrease in slope is likely due to a decrease in interfacial surface area between the electrode and the liquid in the paper compared to the electrode and the bulk solution in the traditional electrochemical cell. Cellulose paper is highly porous and does not fully saturate even when appearing to be fully wetted [61]. Therefore, air will occupy some of the space between the electrode and the paper fluidic. Air does not provide an efficient medium for electron transfer between the solution and the electrode, leading to a smaller interfacial area and a lower measured current [53], [59], [62].

**Figure 3.**
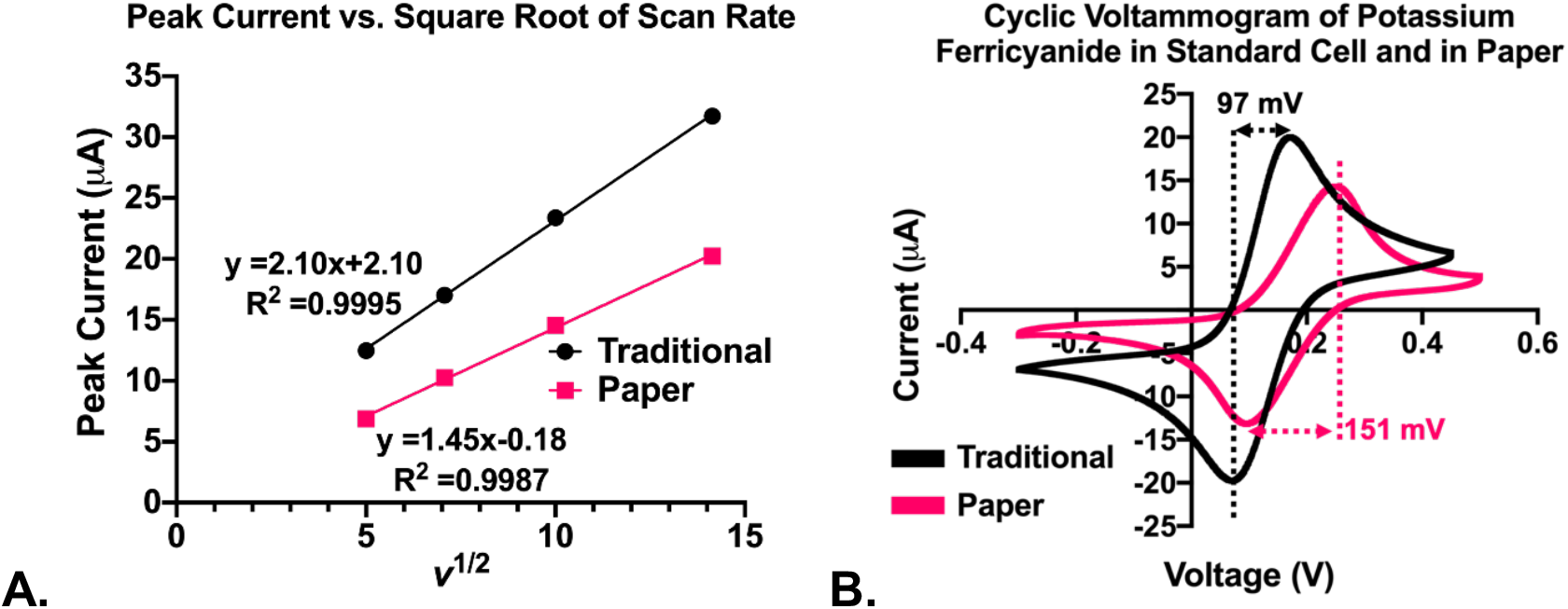
Comparison of potassium ferricyanide sensing in traditional and paper electrochemical cells. **(A)** shows the linear relationship between peak current and the square root of the scan rate (*v*^*1/2*^) in both the traditional electrochemical cell (black) and in paper (pink) as expected by a diffusion-limited reaction. **(B)** shows the difference in peak separation of a reversible electrochemical process with greater peak separation in paper indicating some inhibition of electron transfer efficiency relative to the traditional electrochemical cell. The scan rate was 100 mV/s in figure B. Solutions were kept at room temperature, a pH of 7.4, and measurements were referenced to an Ag/AgCl electrode.

The results shown in Figure 3B indicate redox peak separations of 151.2 mV and 97.1 mV, in the paper and traditional cells respectively. In an ideal reversible electrochemical reaction, the peak separation should be 59 mV for a single electron transfer reaction [53]. The separation greater than 59 mV between the reduction and the oxidation peaks indicates some deviation from the ideal response. However, the ideal response of 59 mV is very rarely observed in practice [53], [60]. Other researchers have reported peak separations as high as 270 mV with potassium ferricyanide in a traditional electrochemical cell [53], [60]. This may occur because molecules adsorb to the surface of the electrode during the experiment, causing a decrease in charge transfer efficiency and rate of the electron transfer, widening the redox peaks [53], [60]. In accordance with these literature findings, we concluded that the redox peak separations of 151.2 and 97.1 mV indicate that paper does not substantially impede the electron transfer efficiency and is a suitable platform for our electrochemical analysis.

### Evaporation Rate of Water from Chromatography Paper and Absorption Capacity

The paper fluidic device drives fluid flow using evaporation at physiological sweat rates. Sweat rates can range from a low of 0.1 *μ*L/min/cm^2^ to a high of 10-20 *μ*L/min/cm^2^ during high intensity workouts [29], [63], [64]. We quantified the evaporation rate of water from the paper, as seen in Figure 4. Evaporation rates from the paper ranged from a high of 0.5 *μ*L/min/cm^2^ (+/-0.05 *μ*L/min/cm^2^) at a humidity of 37% (+/-0.90%) to a low of 0.3 *μ*L/min/cm^2^ (+/-0.01 *μ*L/min/cm^2^) at a humidity of 56% (+/-4.2%) (Figure 4). Our results demonstrate an expected humidity-dependent evaporation rate. Therefore, we will be able to dynamically tune our flow rate based on the variable humidity of the laboratory environment to match that of human sweat rates.

**Figure 4.**
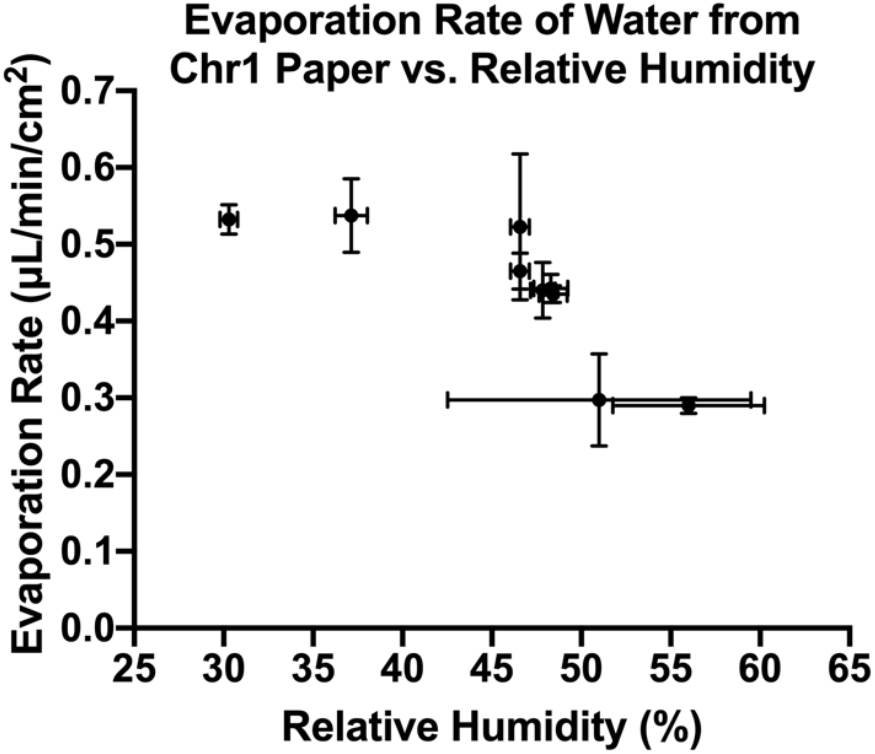
Evaporation rate of water from the paper as a function of relative humidity. Error bars represent standard deviation. N = 2 for the humidity measurements and N = 5 for the evaporation rates. Two humidity measurements were taken. One measurement was taken at the start of the 30-minute evaporation period, and one taken at the end. Evaporation measurements represent the average and standard deviation across 5 pieces of chromatography paper measured on 9 different days.

To ensure saturation of the paper fluidic device, we determined the paper had a water absorption capacity of 34 *μ*L/cm^2^ (+/-11 *μ*L/cm^2^). This allows us to calculate that 162 *μ*L of water is needed to saturate our device with its total surface area of 4.75 cm^2^. As noted in the characterization section, fluctuations in saturation would change the interfacial area between the electrode and the fluid. Variable interfacial area would affect the measured Faradaic current independent of the concentration of cocaine [53], confounding our results. Therefore, it is important to calculate the volume of fluid required to maintain consistent saturation in the device during our continuous, under flow experiments given the variable humidity within the laboratory environment.

### Multi-Hour Cocaine and Buffer Exchange Measurements in Paper Fluidic Device

The paper fluidic device was designed to simulate a fully wearable, continuous, real-time sweat biosensor by measuring cocaine and PBS under flow at physiological sweat rates. Figure 5 demonstrates the time evolution in the Faradaic current measured by the benchtop potentiostat (Figure 5A) and KickStat (Figure 5B) as 0.5 mM cocaine flowed past the sensor, was replaced by PBS, and vice versa. The signal reaches a local maxima and minima corresponding to peak saturation by cocaine and PBS, respectively. The paper was first primed with 0.5 mM cocaine at time, t<0, then PBS was pumped into the system until time, t1. From t1 to t2, cocaine was pumped into the system, then PBS from t2 to t3, cocaine from t3 to t4, and PBS after t4. The signal gain with cocaine, measured by the benchtop instrument (Figure 5A), is roughly 25-37% relative to the subsequent PBS baseline, reaching maximums of 1.26 µA, 1.22 µA, and 1.17 µA in cocaine compared to minimums of 0.92 µA, 0.93 µA, and 0.94 µA in PBS.

**Figure 5.**
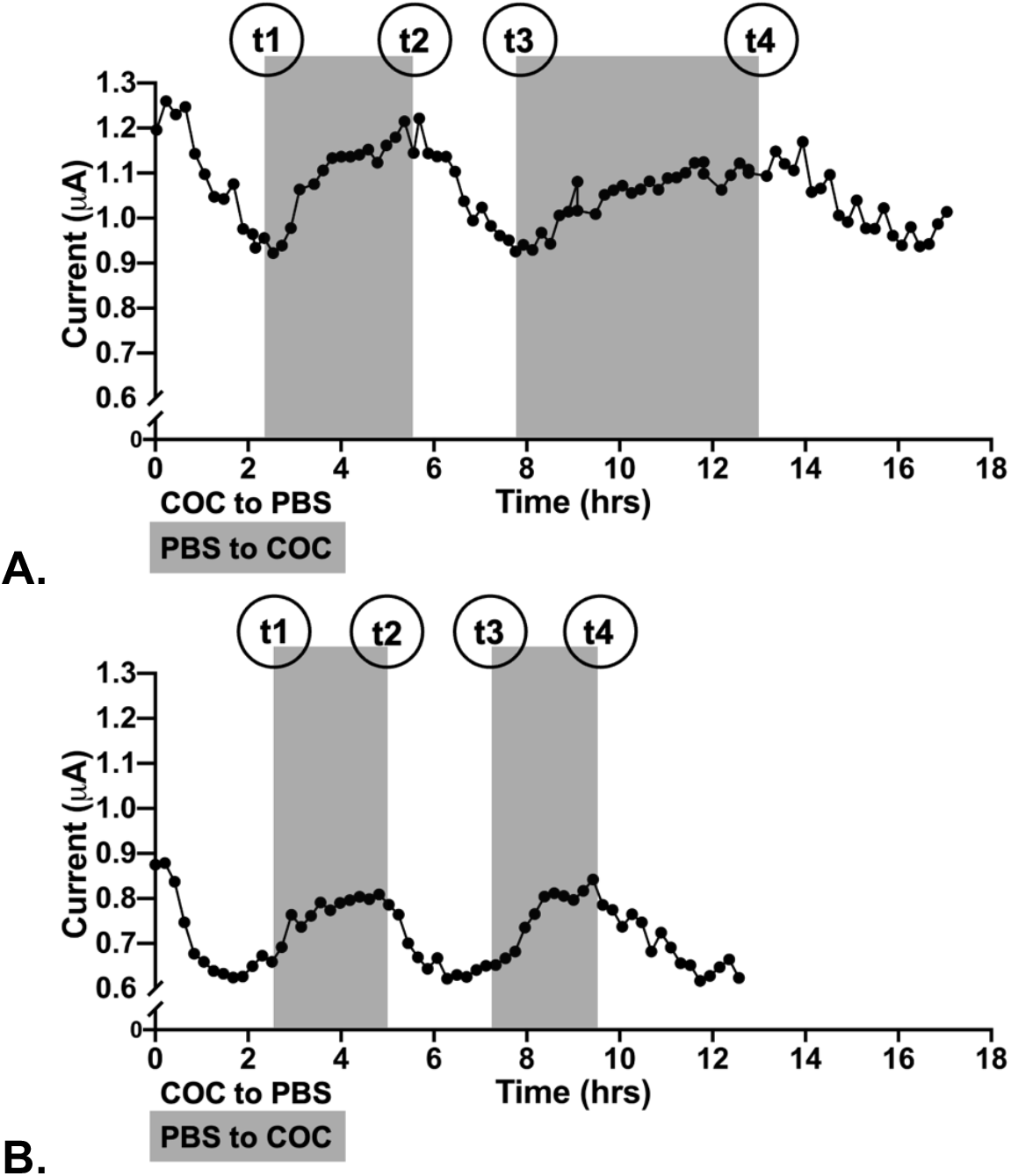
Time evolution of faradaic current measured from cocaine-binding aptamer, in paper under evaporation-driven flow using benchtop (A) and KickStat (B) instruments. Paper was primed with 0.5 mM cocaine (t<0), then PBS was pumped into the system (t>0). PBS was switched to 0.5 mM cocaine at t1, cocaine was switched to PBS at t2, PBS was switched to cocaine at t3, and cocaine was switched to PBS at t4. Solutions were kept at room temperature, a pH of 7.4, and measurements were referenced to an Ag/AgCl electrode.

The currents measured with KickStat (Figure 5B) are smaller than the currents measured with the benchtop instrument, reaching maximums of 0.88 µA, 0.81 µA, and 0.84 µA and minimums of 0.63 µA and 0.62 µA twice. The discrepancy in current between the two datasets may be the result of expected variation in the resulting monolayers of the prepared electrodes [48], [52]. KickStat was previously validated against the benchtop potentiostat in an earlier study and showed no significant disagreement [50].

KickStat measured a signal gain with cocaine of roughly 30-41% relative to the subsequent PBS baseline, while the benchtop potentiostat reported a signal gain of 25-37%. Previous reports using this cocaine aptamer sequence in a traditional electrochemical cell demonstrated a signal gain of 31.4% in 0.44 mM cocaine [21]. The 25-37% signal gain in our paper fluidic device measured by the benchtop instrument and 30-41% measured by KickStat are within the range of the previous literature, indicating measurements of cocaine in the paper fluidic device are accurate and reliable. It is interesting to note that KickStat measured higher percent signal gains than the benchtop potentiostat.

Despite variable temperatures and humidity in the research lab setting, evaporative-driven flow in paper successfully allowed for the aptamer-based detection of cocaine in PBS over the course of 18 hours at fluid delivery rate of 0.3-0.5 *μ*L/min/cm^2^. This rate is well within the physiologic sweating rates of 0.1 *μ*L/min/cm^2^ to a high of 10-20 *μ*L/min/cm^2^ during high intensity workouts [29], [63], [64]. However, under dry conditions or low temperature or low-activity effort, the sweat rates may be even lower than 0.1 *μ*L/min/cm^2^. In such circumstances, sweat induction by pilocarpine [65] or iontophoresis [35] could be used to produce a sweat rate that ensures sensor hydration.

Given the length of the multi-hour experiments, significant baseline drift is expected [55]. Indeed, we observe drift in our experiment as the signal never fully returns to its previous maxima or minima. Nevertheless, we still observe marked increase in signal after 12+ hours of continuous use while also being exposed to ambient laboratory conditions. Future work could correct for baseline drift by including a reference sensor e.g., with a non-binding aptamer, to quantify changes in drift over an extended experiment period. Potentially, differential measurements of voltammetric signals at different pulse frequencies may also improve future signal-to-noise ratios [56], [57].

The physiologically relevant concentrations of cocaine hydrochloride in sweat are closer to 10 nM [15], [28], [30], [63], which is much lower than the concentrations tested here. Nevertheless, the feasibility of detecting cocaine in paper under flow at physiological sweat rates was demonstrated herein. We chose this cocaine aptamer because it is well-studied in the literature [21], [23], [66] and we have extensive previous experience with the sensor [50]. We anticipate that improved conjugation techniques and new aptamers for cocaine and other small molecules can be easily integrated into our system.

A further note, which is not often highlighted in aptasensor work is that the preparation of the cocaine aptasensors is fairly extensive, requiring 5-8 hours, and the sensors were used immediately for optimal sensitivity. We did not explore sensor storage in this paper, however other groups have shown that cocaine aptamer sensors remains effective after 28 days in dry storage with bovine serum albumin as a stabilizing agent [49].

Finally, we were initially concerned that the paper fibers might strip the aptamer from the electrodes by abrading the electrodes over time. In high movement situations, this might still be a possibility, however, as our test fixture is stationary, we did not observe anything that suggests the electrode is being abraded by the paper over time. The incorporation of flexible electrodes in future work, could further limit the potential for abrasion in this type of wearable sensor platform. Additionally, we anticipate that flexible electrodes would conform better to the paper, further improving the interfacial area between the electrode and the saturated paper, increasing signal output.

## Conclusions

We have demonstrated the ability to detect cocaine in paper, under evaporative-driven flow using an electrochemical sensor and our own custom-designed, miniaturized potentiostat. We have provided a proof-of-concept of a wearable system for monitoring concentrations of cocaine in sweat by using our miniaturized potentiostat in place of a traditional, large benchtop instrument. The sensor functioned over the course of 17 hours, enough for a day’s worth of measurements at physiological flow rates. Our paper fluidic platform demonstrates a flow system driven entirely by evaporation. To our knowledge, evaporation-driven flow has not previously been shown in the literature particularly at physiological flow rates. This work demonstrates the feasibility of a paper fluidic device for detecting cocaine and lays the foundation for real-time detection of illicit drugs in sweat using a miniature wearable device.

## Acknowledgements

This work was supported by the National Institute of Health National Institute of Drug Abuse [contract number 108380] and the Ralph W. and Grace M. Showalter Research Trust Award. We would like to thank Dr. Peter Kissinger, Dr. Frédérique Deiss, Dr. Netz Arroyo, Dr. Rebecca Lai, and Dr. Kevin Plaxco for their advice on electrochemistry techniques. Orlando S. Hoilett was supported by the Geddes Fellowship.

